# Human neural rosettes secrete bioactive extracellular vesicles enriched in neuronal and glial cellular components

**DOI:** 10.1101/2024.01.10.574187

**Authors:** Malena Herrera Lopez, Matías Bertone Arolfo, Mónica Remedi, Laura Gastaldi, Carlos Wilson, Gonzalo G. Guendulain, Danilo Ceschin, Andrés Cardozo Gizzi, Alfredo Cáceres, Ana Lis Moyano

## Abstract

Extracellular vesicles (EVs) play a critical role in the development of neural cells in the central nervous system (CNS). Human neural rosettes (hNRs) are radial cell structures that assemble from induced pluripotent stem cells (hiPSCs) and recapitulate some stages of neural tube morphogenesis. Here we show that hiPSCs and hNRs secrete EVs (hiPSC-EVs and hNR-EVs) with distinct protein cargoes. Remarkably, hNR-EVs carry neuronal and glial cellular components involved in CNS development. By *in silico* analysis, we found hNR-EVs protein signature is expressed *in vivo* and *in vitro* during human brain development. Importantly, hNR-EVs stimulate hiPSCs to change their cellular morphology with a significant reduction in the pluripotency regulator SOX2. Interestingly, these effects were inhibited by antibodies against an unexpected neuroglial cargo of hNR-EVs: the major proteolipid protein (PLP1). These findings show that hNRs secrete bioactive EVs containing neural components and might contribute as trophic factors during human CNS development.

## Introduction

Human neural rosettes (hNRs) are an assembly of neural cells generated *in vitro* from induced pluripotent stem cells (hiPSCs). These radial structures exhibit apicobasal polarity, resembling the cellular architecture of the human neural tube. hNRs generate neurons and glial cells of the central nervous system (CNS) recapitulating key molecular and biological events associated with human brain morphogenesis and development (Conti and Cattaneo, 2010; Elkabetz and Studer, 2008). hNRs exhibit a heterogeneous population of cells including neuroepithelial, neural stem and progenitor cells that potentially can differentiate into any CNS cell type. Therefore, hNRs are a promising *in vitro* model to study human CNS development (Mertens et al., 2016).

Several trophic factors are used to maintain hNRs *in vitro* but little is known about the secreted cellular components that may contribute to spatiotemporal coordination of hNRs formation. Extracellular vesicles (EVs) are nanosized vesicles that transport lipids, proteins and nucleic acids among different CNS cell types and their cargoes can elicit a phenotypic response in acceptor cells (Holm et al., 2018; Van Niel et al., 2018). EVs secreted in CNS development might regulate proliferation and differentiation of stem and progenitor cells essential for neural growth and localization (Bahram Sangani et al., 2021; Schnatz et al., 2021). However, little is known about EVs biology, their cargoes and biological significance during fetal development of the human CNS.

Neuronal and glial cellular components are secreted in EVs from different animal models and human stem cells. The myelin proteolipid protein (PLP1) and its spliced isoform DM20 are secreted by oligodendrocytes (glial cells that synthetize and assemble CNS myelin), also detected in EVs isolated from rodent brains (Frühbeis et al., 2020; Krämer-Albers et al., 2007). Although PLP1 is the most abundant protein in CNS myelin in postnatal and adult brains, it is also expressed during embryonic stages, *i.e.* long before myelin is synthetized and assembled (Delaunay et al., 2008; Spassky et al., 2000; Timsit et al., 1995). Currently, little is known about the association between PLP1 and EVs in humans, as well as its spatiotemporal expression pattern during early neurodevelopment (Kronquist et al., 1987).

Here we show for the first time that hiPSCs and hNRs secrete EVs (hiPSC-EVs and hNR-EVs, respectively) enriched in proteins associated with EVs. Only hNR-EVs are specifically enriched in cellular components involved in CNS development, usually expressed at different developmental stages *in vitro* and *in vivo*. Interestingly, hiPSCs treated with hNR-EVs exhibit changes on their morphology and a significant reduction of protein levels of SOX2, a well-established molecular marker of pluripotency. Remarkably, these effects were inhibited by antibodies against PLP1, an unexpected cargo of hNR-EVs. In conclusion, our data suggests that hNR-EVs along with their molecular cargoes might participate as trophic factors or effectors during early human neurodevelopment.

## Materials and Methods

### hiPSCs cell culture and differentiation into hNRs

2 hiPSCs clones (F2A112 and F2A121) from a healthy donor were obtained from PLACEMA Foundation where they were reprogrammed and characterized (Casalia et al., 2021). To obtain hNRs we used a protocol previously described (Zhang and Zhang, 2010), with modifications. Briefly, hiPSCs were cultured on a layer of irradiated mouse (Knockout serum replacement-KSR) in the presence of a ROCK inhibitor (Y-27632) and the fibroblast growth factor (FGF). hiPSCs colonies were manually selected and enzymatically dissociated to be differentiated into embryoid bodies (EB) in medium with Y-27632 and low FGF (4 ng/ml). Between 7-8 days in culture, media was supplemented with N2 and FGF (20 ng/ml) and on day 10 EB were adhered to the surface covered with Geltrex (Invitrogen). After 4-5 days in culture, the area and thickness of the neural rosettes increased (definitive neuroepithelium). Cell cultures in all stages were grown in defined culture medium to rule out the incorporation of EVs present in media supplemented with sera (Wiklander et al., 2015). hiPSCs- and hNR-conditioned media were collected every other day.

### Immunocytochemistry and confocal microscopy

Cells were fixed in 4% paraformaldehyde (PFA) at room temperature (RT) for 15 minutes and then washed 3 times with phosphate buffered saline (PBS). Then, they were permeabilized with PBS-Triton 0.25% and blocked for 1 hour with bovine serum albumin 5% in PBS-Triton 0.1% at RT. Subsequently, some of the following primary antibodies were incubated 24 hours at 4 °C: rabbit anti-SOX2 (1:100 dilution, Abcam, ab137285), rabbit anti-doublecortin (DCX, 1:1000 dilution, Abcam, ab207175), mouse anti-tyrosinated tubulin (TUB-1A2, 1:1000 dilution, Sigma, T9028), chicken anti-nestin (1:500 dilution, Abcam, ab134017) and rat AA3 monoclonal hybridoma anti-proteolipid protein 1 (PLP1, 1:100 dilution). AA3 was a kind gift of Dr. Irene Givogri and Dr. Ernesto Bongarzone. Next, the following secondary antibodies were incubated at RT for 1h: anti-IgG rabbit (1:1000; Alexa Fluor 488 or 546, Life Technologies, A11008 or A11010), anti-IgG mouse (1:1000 dilution, Alexa Fluor 546; Life Technologies, A11030), anti-IgY chicken (1:1000 dilution, Alexa Fluor 488; Invitrogen, A11039) and/or anti-IgG rat (dilution 1:1000, Alexa Fluor 568; Molecular Probes, A11077). Cell nuclei were labeled with dapi (4 ’, 6-diamidino-2-phenylindole, Invitrogen, P36931). Images were obtained in the Zeiss LSM 800 Confocal Optical Microscope at Centro de Micro y Nanoscopía de Córdoba (CEMINCO-CONICET-UNC) and analyzed using ImageJ image (NIH, Bethesda, MD).

### Isolation of hiPSC-EVs and hNR-EVs

EVs were isolated as described previously (Moyano et al., 2016; Thery et al., 2006). Briefly, cell culture conditioned media was collected from hiPSCs and hNRs (2-10 ml) and centrifuged at low speed (300 xg) to remove large particles. Supernatants were centrifuged 10 minutes at 2000 xg and then this supernatant filtered through a 0.22 µm filter. EVs were isolated after ultracentrifugation at 100,000 xg for 90 minutes. Isolated EVs were washed in 10 ml of PBS and pelleted again at 100,000 xg for 90 minutes. EVs were resuspended in 50 µl of 0.22-µm-filtered PBS. All centrifugation and ultracentrifugation steps were performed at 4 °C. hiPSC-EVs and hNR-EVs samples were stored at -80 °C until further analysis.

### Treatment of hiPSCs with hNR-EVs

Small hiPSCs colonies were cultured in coverslips on a layer of MEFi in serum-free medium for 24 h before treatment. Cells were treated with hNR-EVs isolated from 0.5 ml hNR-conditioned media in a final volume of 0.5 ml serum-free medium and incubated 24 h at 37 °C. Antibody-mediated inhibition of hNR-EVs was performed with a commercial antibody that recognizes an extracellular topological domain of PLP1 at its N terminus (AA 36 to 85, Invitrogen, PA5-40788) incubating hNR-EVs with anti-PLP1 1.25 ug/ml for 15 min. before hiPSCs treatment. After treatment cells were fixed in 4% PFA for immunocytochemistry. Two biological replicates per clone (F2A112 and F2A121) were used for hiPSCs and hNR-EVs.

### Electron microscopy and dynamic light scattering

hiPSC-EVs and hNR-EVs were fixed in 2% PFA and characterized by transmission electron microscopy (TEM) at the Centro de Microscopía Electrónica (INTA-CIAP) using the TEM Jeol 1200 EX II 14.33 electron microscope as previously described (Thery et al., 2006). Particle size distribution was analyzed by dynamic light scattering (DLS) using SZ-100 nanopartica series instruments (Horiba) at the Centro de Investigaciones en Química Biológica de Córdoba (CIQUIBIC-CONICET-UNC).

### Protein quantification

Proteins were measured by the bicinchoninic acid (BCA, Pierce, 23227) method. Using the EL800 BioTek microplate reader to determine the absorbance at 562 nm.

### LC-MS analysis

Peptide separations were carried out on a nanoHPLC Ultimate3000 (Thermo Scientific) using a nano column EASY-Spray ES901 (15 cm × 50 μm ID, PepMap RSLC C18). The mobile phase flow rate was 300 nl/min using 0.1% formic acid in water (solvent A) and 0.1% formic acid and 100% acetonitrile (solvent B). The gradient profile was set as follows: 4-35% solvent B for 30 min, 35%-90% solvent B for 1 min and 90% solvent B for 5 min. Two microliters of each sample were injected. MS analysis was performed using a Q-Exactive HF mass spectrometer (Thermo Scientific). For ionization, 1,9 kV of liquid junction voltage and 250 °C capillary temperature was used. The full scan method employed a m/z 375– 1600 mass selection, an Orbitrap resolution of 120000 (at m/z 200), a target automatic gain control (AGC) value of 3e6, and maximum injection times of 100 ms. After the survey scan, the 20 most intense precursor ions were selected for MS/MS fragmentation. Fragmentation was performed with a normalized collision energy of 27 eV and MS/MS scans were acquired with a dynamic first mass, AGC target was 5e5, resolution of 30000 (at m/z 200), intensity threshold of 4.0e4, isolation window of 1.4 m/z units and maximum IT was 200 ms. Charge state screening enabled to reject unassigned, singly charged, and equal or more than seven protonated ions. A dynamic exclusion time of 15s was used to discriminate against previously selected ions. LC-MS analysis was performed at the Unidad de Espectrometría de Masa, Instituto de Biología Molecular y Celular de Rosario (UEM-IBR-CONICET).

### MS data analysis

MS data were analyzed with Proteome Discoverer (version 2.4) (Thermo) using standardized workflows. Mass spectra *.raw files were searched against the database of Homo sapiens from Uniprot (UP000005640) Precursor and fragment mass tolerance were set to 10 ppm and 0.02 Da, respectively, allowing 2 missed cleavages, carbamidomethylation of cysteines as a fixed modification, methionine oxidation and acetylation N-terminal as a variable modification. Identified peptides were filtered using Percolator algorithm with a q-value threshold of 0.01.

### GO enrichment analysis

MS datasets were used as input for GO enrichment analysis to detect potential molecular function, biological processes and cellular components GO terms using the g:Profiler2 (version e107_eg54_p17_bf42210) (Raudvere et al., 2019). GO terms with g:SCS multiple testing correction method and a significance threshold of 0.05 were considered enriched. Enriched GO terms were visualized using Microsoft Excel (version 2212). GO subsets for EV-related proteins were selected with the following terms: extracellular vesicle, vesicle, vesicular, exosome and exosomal. GO subsets for neuronal- and glial-related proteins were selected with the following terms: neural, neuron, axon, glia, astrocyte, microglia, Schwann, oligodendrocyte, myelin, nervous, nerve, brain. Top 100 EVs proteins were fused from Vesiclepedia (Kalra et al., 2012) and Venn diagrams were created with https://molbiotools.com/listcompare.php).

### In silico analyses

Gene Expression Omnibus (GEO) database freely archives and distributes public microarray, next-generation sequencing (NGS) results, and other forms of genomic data that can be combined and reanalyzed to reveal previously unknown relationships (Edgar, 2002). Using GEO repository, we analyze RNAseq datasets from 3D hiPSC-derived neural spheroids (GEO accession: GSE102139 from Simão et al., 2018) and hNRs derived from hESCs (GEO accession: GSE65369 from Edri et al., 2015). To analyze scRNAseq datasets we applied UCSC Cell Browser (Speir et al., 2021) with datasets from hiPSC-derived brain organoids (GEO accession: GSE124299 from Pollen et al., 2019) and human fetal brain (Data available at dbGaP: phs000989.v3 from Nowakowski et al., 2017). The MS datasets were analyzed from PRIDE (Perez-riverol et al., 2022) datasets of cerebroids (Accession number: PXD011605 from Nascimento et al., 2019) and human fetal brain (Accession number: PXD004076 from Djuric et al., 2017).

### Western Blot

Samples were lysed with RIPA buffer (50 mM Tris-HCl pH 8, 1% v / v triton X-100, 1mm EDTA and 0.15M NaCl) and the protein concentration determined by BCA. The samples were diluted in Laemmli buffer (0.25 % w / v Bromophenol Blue, 15 % v / v b-Mercaptoethanol, 50 % v / v Glycerol, 10 % w / v SDS and 0.25 M Tris-HCl pH 6 , 8) and incubated at 95 ° C for 10 minutes. Lysates (5-10 µg protein) were resolved by sodium dodecyl sulfate 12% polyacrylamide gel electrophoresis (SDS-PAGE), at 200 V for 1 hour (BioRad). Gels were transferred to a nitrocellulose or PVDF membrane at 100 V for 90 minutes at 4 °C. These membranes were incubated with a blocking solution (5% w / v non-skim milk diluted in TBS + 0.05% v / v Tween) for 1 hour. After, incubated for 24 hours at 4 ° C with the following primary antibodies: rabbit anti-syntenin 1 (1:2000 dilution, Abcam, ab133267) and mouse anti-tubulin α (1:2000 dilution, clone DM1A, Sigma, T9026). Membranes were washed 3 times with PBS-Tween 0.01% and incubated with peroxidase-labeled secondary antibodies to detect antibody reactivity by enhanced chemiluminescence (ECL).

### Statistical analysis

Data were analyzed using Student’s t-test or one-way ANOVA followed by Dunnett’s or Tukey’s multiple comparison tests. P values <0.05 were considered significant. Data was examined and visualized using Microsoft Excel (version 2212), the GraphPad Prism 10.00 program (San Diego, California, www.graphpad.com).

## Results

### Cell culture model and EVs characterization

Human induced pluripotent stem cells (hiPSCs) were differentiated into neural rosettes (hNRs) and characterized by immunofluorescence (Figure 1A and 1B). hiPSCs colonies displayed their typical morphology expressing the pluripotency marker SOX2. hNRs showed radially-organized cells expressing SOX2 and the neural stem cells marker nestin. EVs were isolated by ultracentrifugation from serum-free conditioned media from hiPSCs and hNRs (Figure 1C). EVś morphology and size distribution were evaluated using transmission electron microscopy (TEM) and dynamic light scattering (DLS). By TEM and DLS we found that hiPSC-EVs and hNR-EVs exhibit their typical morphology and size distribution (Figure 1D and 1E). The most abundant EVs showed a mean size of 378.9 nm (79.3 % EVs) in hiPSC-EVs and 188.6 nm (92.1 % EVs) in hNR-EVs. These results showed that hiPSCs and hNRs secrete EVs with a heterogeneous size distribution.

**Figure 1.**
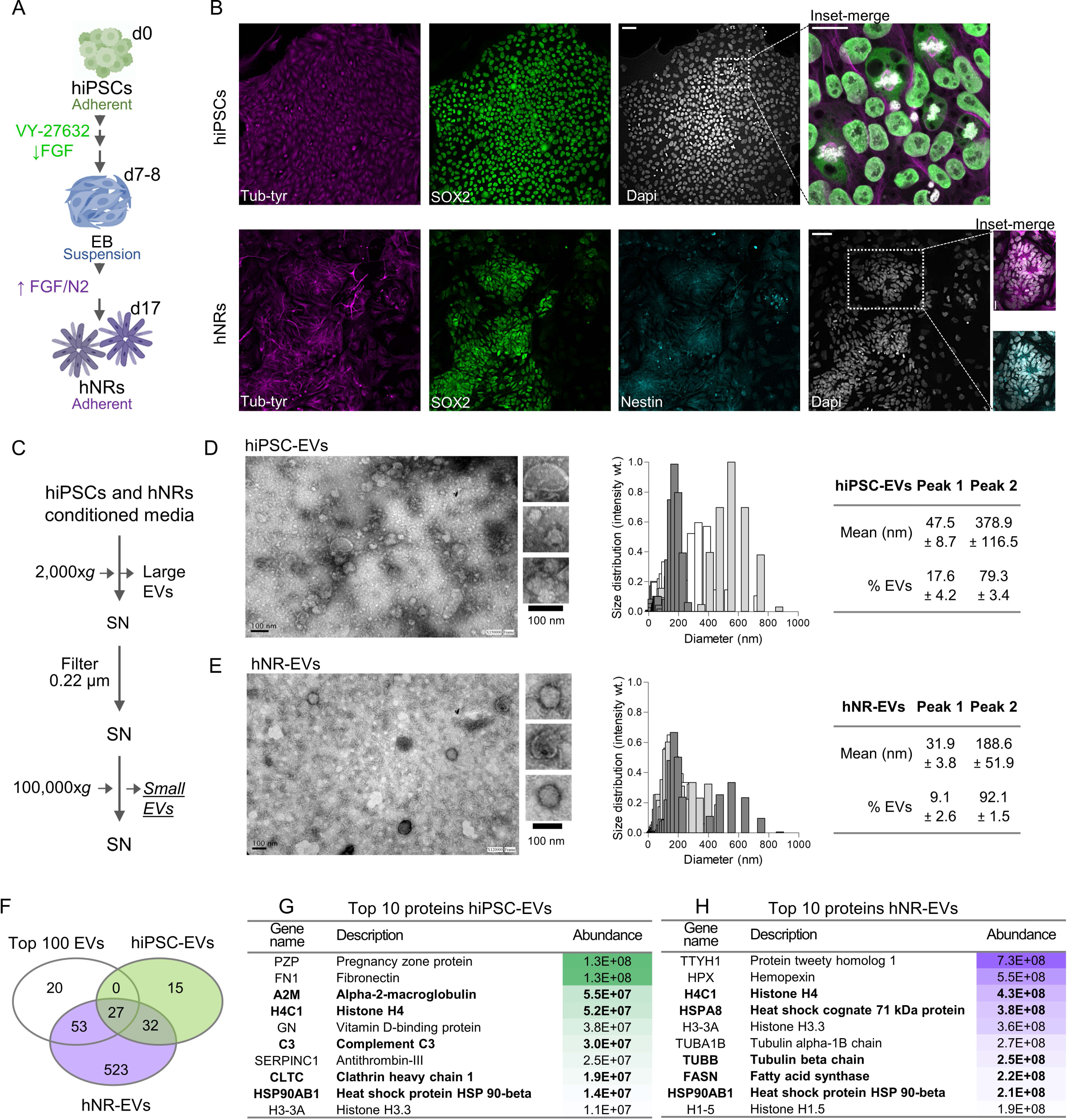
hiPSCs and hNRs cell culture. (A) hiPSCs cultures and differentiation into hNRs to obtain cell culture conditioned media. (B) Confocal micrographs and immunocytochemistry of hiPSCs and hNRs with antibodies against tyrosinated tubulin (Tyr tub, magenta), SOX2 (green) and nestin (cyan). Dapi (grays). Scale bars: hiPSCs 50 μm (10x) and inset 20 μm (63x). hNRs 50 μm (20x) and inset 20 μm (20x). **hiPSC-EVs and hNR-EVs characterization**. (C) Isolation of small EVs by differential ultracentrifugation. SN: supernatant. TEM micrographs and size distribution by DLS of hiPSC-EVs (D) and hNR-EVs (E). Data: mean ± SEM. **hiPSC-EVs and hNR-EVs are enriched in EV-related proteins.** (F) Venn diagram showing the overlap between top 100 EVs proteins (Vesiclepedia) and those identified by MS analysis in hiPSC-EVs (light green) and hNR-EVs (light purple). Top 10 proteins identified in hiPSC-EVs (G) and hNR-EVs (H). Highlighted proteins are common to top 100 EVs, hiPSC-EVs and hNR-EVs.

### hiPSC-EVs and hNR-EVs are associated with EVs markers

To further characterize hiPSC-EVs and hNR-EVs we analyzed their protein composition by mass spectrometry (MS). hiPSC-EVs showed significantly higher levels of total protein content compared with hNR-EVs (Figure S1A) and MS analysis identified 74 proteins in hiPSC-EVs and 635 in hNR-EVs (Table S1). Compared with the top 100 proteins associated with EVs from the database Vesiclepedia (Kalra et al., 2012), we found 27 proteins overlapping with hiPSC-EVs and 80 with hNR-EVs (Figure 1F, S1B and table S1). Some of them are among the top 10 proteins detected in hiPSC-EVs and hNR-EVs (Figure 1G and 1H). Also, both preparations are enriched in MISEV 2018 markers (Kugeratski et al., 2021; Théry et al., 2018) with only apolipoprotein A-1 as contaminant (Figure S1C). Collectively, these results showed that our preparations are enriched in EVs markers and hNR-EVs exhibit a more complex protein signature compared with hiPSC-EVs.

### hNR-EVs are associated with neuronal and glial cellular components

Gene ontology (GO) enrichment analysis with proteins from hiPSC-EVs and hNR-EVs revealed diverse terms among the top 10 significantly enriched in molecular functions (MF), cellular components (CC) and biological processes (BP) (Figure S2). Both preparations showed a significant enrichment in filtered GO terms related to EVs in CC and BP (Figure 2A and 2B). Furthermore, filtered GO terms related to neuronal and glial CC and BP categories were significantly enriched almost exclusively in hNR-EVs (Figure 2C and 2D). Interestingly, hNR-EVs protein signature exhibits unique and overlapping proteins related to GO terms EVs, CNS development and myelin sheath (Figure 2E, table S1). Among common proteins we found ITGB1, CNP, HSP90AA1, CALM3 and unexpectedly PLP1, the most abundant protein in CNS myelin (Figure 2F). These results indicate that both preparations exhibit proteins associated with EVs. Moreover, only hNR-EVs contain proteins related to CNS development and remarkably to myelin sheet (myelination is a postnatal process).

**Figure 2.**
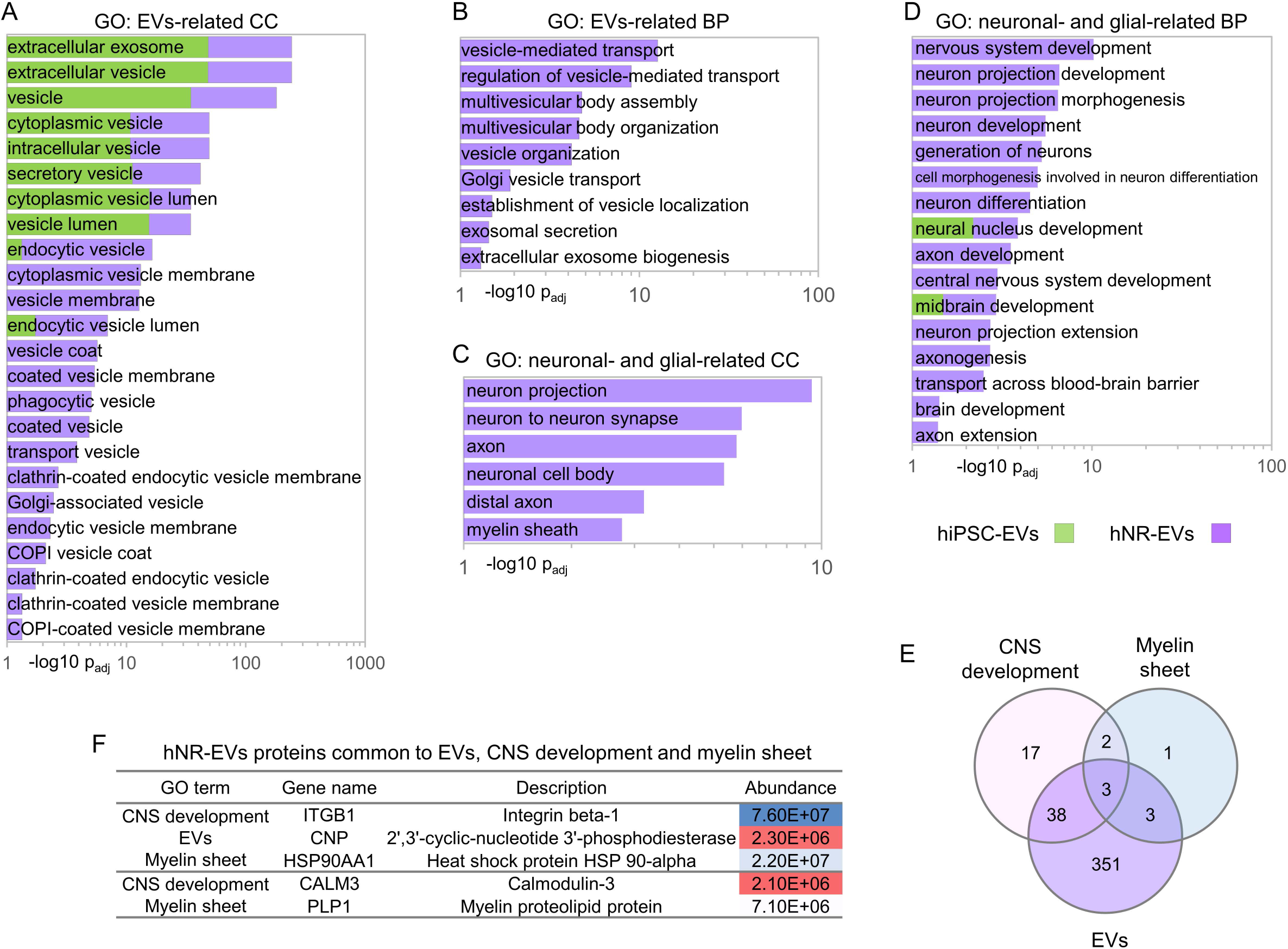
hiPSC-EVs and hNR-EVs are enriched in proteins related to EVs. (A and B) GO analysis showing enrichment (- log10 of adjusted p-value) of GO terms related to EVs proteins in hiPSC-EVs (light green) and hNR-EVs (light purple) between CC and BP groups. **hNR-EVs are enriched in proteins related to neuronal and glial CC and BP**. (C and D) GO analysis showing enrichment of GO terms related to neuronal and glial CC and BP. (E) Venn diagram showing the overlap between proteins identified by MS analysis in hNR-EVs related to GO terms: EVs (GO:1903561, light purple), CNS development (GO:0007417, light red) and myelin sheet (GO:0043209, light blue). (F) Proteins identified by MS analysis in hNR-EVs common to EVs, CNS development and/or myelin sheet (E).

### *In silico* analyses: hNR-EVs proteins are expressed *in vitro* and *in vivo* during development

To further confirm that hNRs express ITGB1, HSP90AA1 and PLP1 under different culture conditions we analyze RNAseq datasets from 3D hiPSC-derived neural spheroids (Simão et al., 2018) and hNRs derived from human embryonic stem cells (hESCs) (Edri et al., 2015). We found that ITGB1, HSP90AA1 and PLP1 are expressed at different time points, culture conditions and cell types (Figure 3A and 3B). Moreover, scRNAseq datasets using UCSC Cell Browser (Speir et al., 2021) showed that their transcripts are expressed in hiPSC-derived brain organoids (Pollen et al., 2019) and human fetal brain (Nowakowski et al., 2017) (Figure 3C). Similar results were observed at the protein level in MS datasets (Djuric et al., 2017; Nascimento et al., 2019) (Figure 3D and 3E). These observations indicate that proteins associated with hNR-EVs related to CNS development are expressed at various time points during human fetal brain development *in vivo* and *in vitro* before myelination.

**Figure 3.**
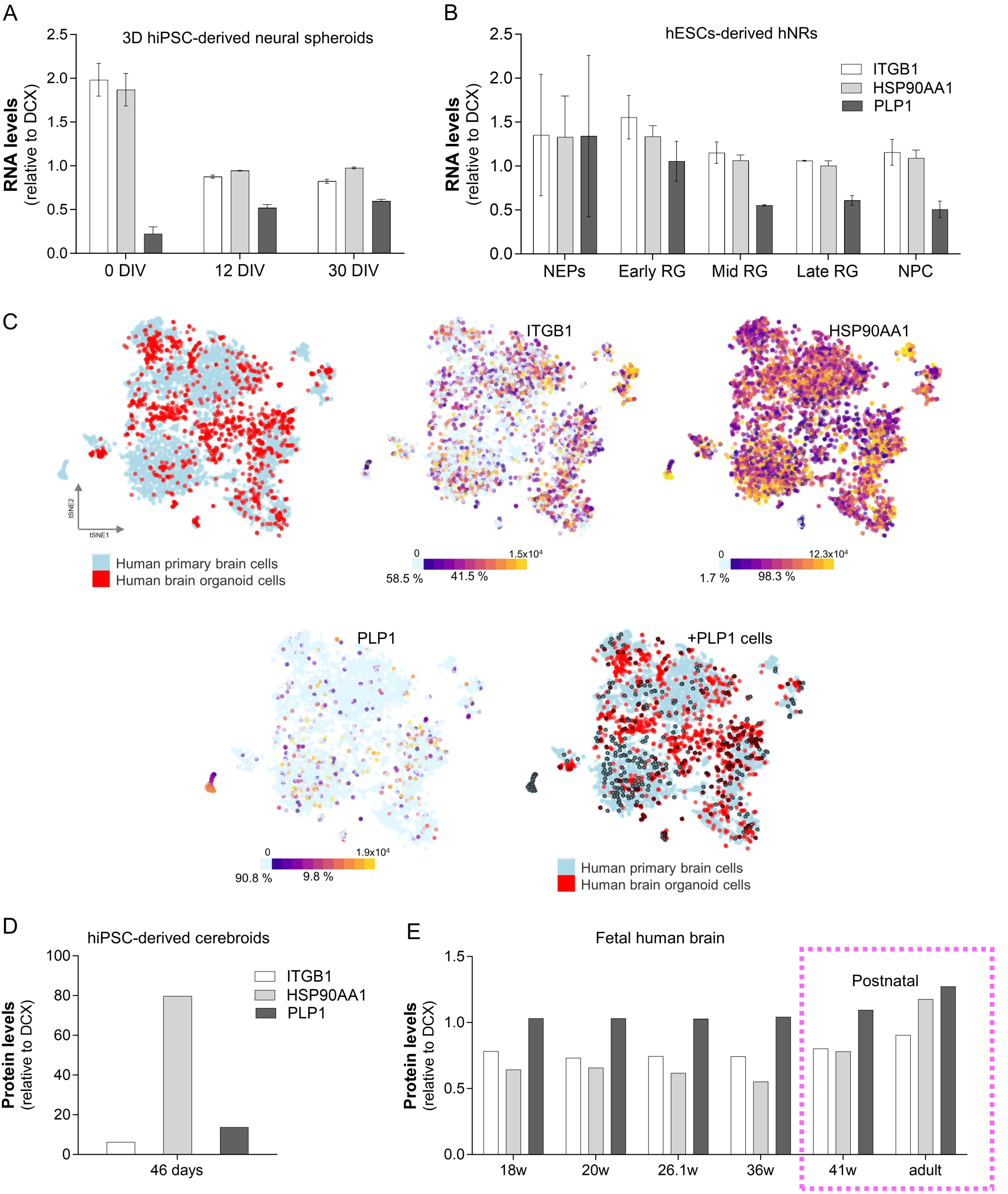
hNR-EVs proteins in hNRs, cerebroids and human brain. Expression of ITGB1, HSP90AA1 and PLP1 in RNAseq datasets from (A) 3D hiPSC-derived neural spheroids and (B) hNRs derived from hESCs. NEPs: neuroepithelial, RG: radial glia and NPC: neural progenitor cells relative to DCX levels (immature neuron marker). (C) tSNE plots highlighting the expression of ITGB1, HSP90AA1 and PLP1. Frequency of cells in scRNAseq datasets from human primary (fetal) brain cells (light blue) and cerebroids (red). +PLP1 cells selected (black circles) among human primary brain cells and cerebroids. Protein levels of ITGB1, HSP90AA1 and PLP1 relative to DCX in (D) 46 days cerebroids and (E) human brain: fetal and postnatal.

### Cellular localization of PLP1 and biological activity of hNR-EVs

To validate the expression of PLP1 in hNRs we analyzed by immunofluorescence its cellular distribution in hiPSCs and hNRs. We found that PLP1 exhibits a radial and differential localization in hNRs outward from their central lumen (Figure 4A-4D) but is not expressed in hiPSCs (Figure S2C). To assess hNR-EVs biological activity, we supplemented hiPSCs cultures with hNR-EVs and after EVs treatment 20.8 % hiPSCs switch into +PLP1 cells with a significant decrease in the levels of the marker SOX2 (Figure 4E-4G). We also examined whether antibodies that recognize an extracellular topological domain of PLP1 (amino acids 36 to 85) inhibit hNR-EVs bioactivity (Yamada et al., 1999). We found that anti-PLP1 antibodies significantly reduced +PLP1 cells and restored SOX2 levels (Figure 4F and 4G). Collectively, these findings suggest that PLP1 is expressed in hNR-EVs and induces changes in hiPSCs homeostasis.

**Figure 4.**
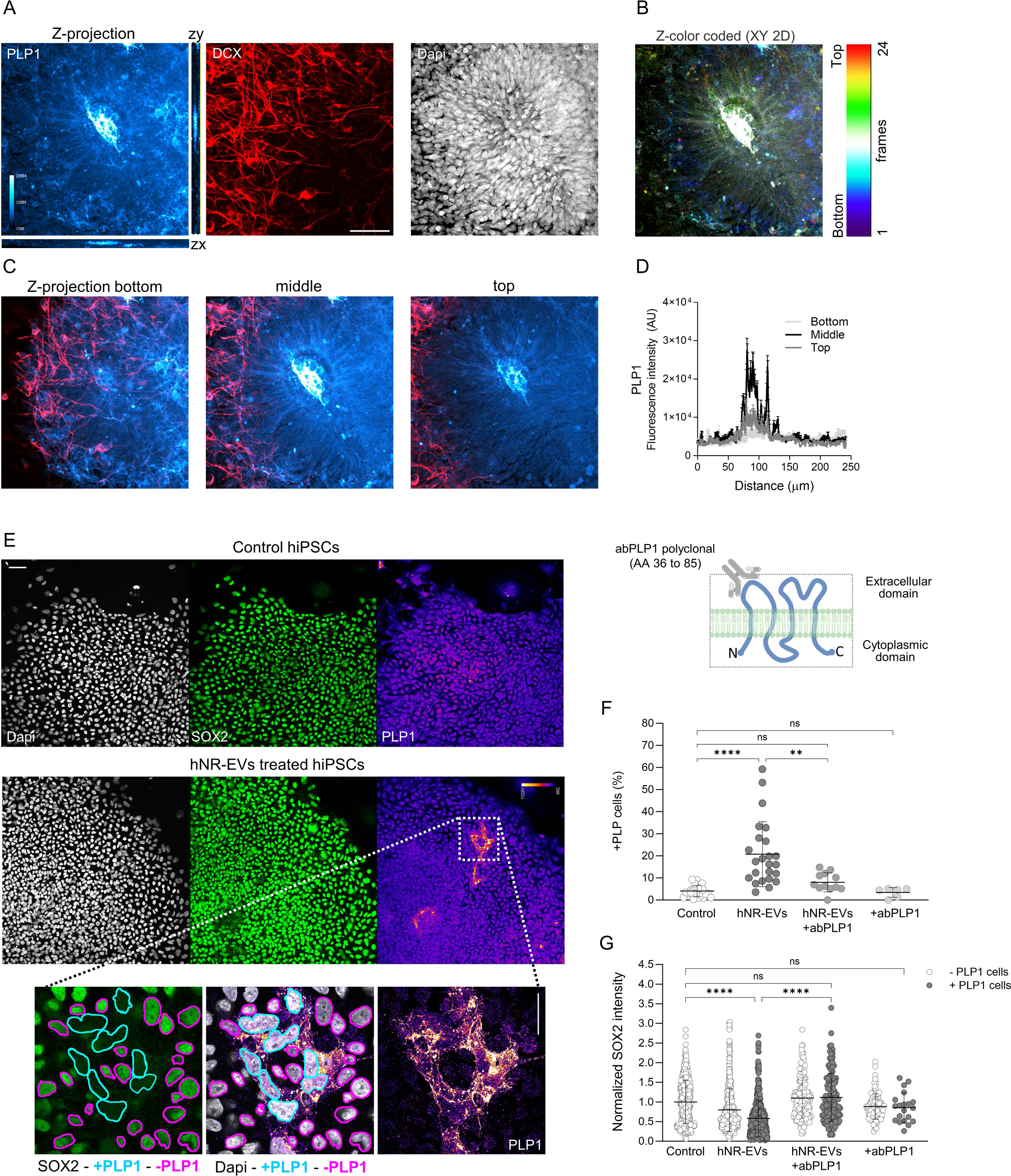
PLP1 has a differential cellular localization in hNRs. (A) Confocal micrographs of hNRs and immunocytochemistry with antibodies to PLP1 (Cyan hot) and DCX (red). Images show maximum projections of confocal Z-stacks and lateral sections. (B) Z-color coded XY 2D image projections of confocal Z-stacks and (C) transversal sections from bottom, middle and to top segments. (D) Fluorescence intensity (AU) in transversal sections. Scale bar: 50 μm at 20x. **Biological activity of hNR-EVs in hiPSCs**. (E) Confocal micrographs and immunocytochemistry of hNRs with antibodies to PLP1 (Fire LUT) and SOX2 (green) in hiPSCs control and treated with hNR-EVs. Scale bar: 50 μm at 20x. Inset from (E) and segmentation: -PLP1 (purple selection) and +PLP1 (cyan selection) cells. Scale bar 25 μm at 63x. Dapi: grays. (F) Percentage of +PLP1 cells per condition. (G) Normalized SOX2 fluorescence intensity in -PLP1 and +PLP1 cells under control or treated conditions. Data: mean ± SD. One-way ANOVA, followed by Tukey’s post hoc test. Ns: no significant and **p < 0.01, ****p < 0.0001 (2 clones: F2A112 and F2A121, 5-6 experiments).

## Discussion

Cell culture models derived by the *in vitro* differentiation of hiPSCs can recapitulate *in vivo* human neural tube formation. Emerging evidence indicates that EVs are key players in the cellular and molecular landscape that emerges during human CNS development. Our study reveals that hiPSCs and hNRs secrete EVs with different protein cargoes. hNR-EVs are enriched in neuronal and glial proteins involved in CNS development expressed at different time points and cell types *in vitro* and *in vivo*. Remarkably, hNR-EVs stimulate hiPSCs to change their cellular morphology with a significant reduction in the pluripotency marker SOX2. These effects were inhibited by antibodies against PLP1, a neuroglial cargo in hNR-EVs. In summary, our work indicates that hNRs secrete bioactive EVs containing neuronal and glial components and might participate as trophic factors in early human CNS development.

Several works have focused on EVs secreted by neural stem and progenitor cells due to their potential regenerative effects in neurological disorders (Holm et al., 2018). However, little is known about EVs secreted during human neural tube formation and CNS development (Bahram Sangani et al., 2021). hiPSCs resemble the inner cell mass of the blastocyst and differentiate into hNRs characterized by an ensemble of neuroepithelial, stem and progenitor cells. Our results showed that hNR-EVs exhibit a more complex protein signature than hiPSC-EVs potentially due to the more diverse cellular landscape of hNRs compared with hiPSCs. These findings indicate that proteins involved in CNS development are selective cargoes of hNR-EVs and might be defined by hNRs spatiotemporal cellular and molecular composition.

PLP1 is the most abundant tetraspan transmembrane protein in CNS myelin and is secreted by murine oligodendrocytes in EVs (Frühbeis et al., 2013; Krämer-Albers et al., 2007). Although is not clear what is the biological significance of PLP1 associated with these vesicles, *Plp1*^−/−^ knockout mice exhibit impaired oligodendroglial EVs release and axonal transport (Frühbeis et al., 2020; Schnatz et al., 2021). Moreover, *in vitro* studies showed that at least a fragment of PLP1 can regulate oligodendrocyte proliferation (Yamada et al., 1999). Our study shows that hNR-EVs biological activity is inhibited by anti-PLP1 antibodies. These observations support the hypothesis that neural components associated with hNR-EVs could mediate EVs release and / or function as modulators of fetal CNS development.

PLP1 and its spliced variant DM20 have been detected in embryonic cells capable of differentiating into neural cells, even before myelination (Campagnoni and Skoff, 2001; Delaunay et al., 2008; Timsit et al., 1995). Loss- and gain-of-function studies in animal models and PLP1 mutations associated with Pelizaeus-Merzbacher disease (PMD) highlight the importance of PLP1 in myelin’s axon-supportive function (Nave and Trapp, 2008; Stadelmann et al., 2019). PLP1/DM20 induce the formation of vesicles and multilamellar assemblies that might influence the shedding of EVs (Bizzozero and Howard, 2002; Ruskamo et al., 2022). Our findings suggest that PLP1/DM20 might have a wider biological role during human CNS development beyond myelination and axonal support. Moreover, hNRs could provide a novel model to study the relevance of EVs biology in PMD and other neurodevelopmental disorders.

EVs derived from stem cells can regulate various biological mechanisms under physiological and pathological conditions, but little is known about their biology during human CNS development (Bahram Sangani et al., 2021). Limited availability of human models and limitations in current EVs isolation methods hinder comprehensive studies on their functional properties. Potentially our EVs preparations are contaminated with other coisolated nanoparticles (Théry et al., 2018). Therefore, we cannot exclude the possibility that the biological effects of hNR-EVs on hiPSCs might be mediated by other components present in the hNRs’ secretome. Thus, more comprehensive studies are necessary to establish the biological role of EVs cargoes in CNS development during neural tube formation.

In summary we found that hiPSCs and hNRs secrete EVs with different protein cargoes. hNR-EVs transport cellular components of neuronal and glial origin present at different developmental stages *in vitro* and *in vivo*. Remarkably, hNR-EVs can induce changes in hiPSCs’ phenotype indicating their biological functionality. Future studies are needed to address what is the biological role of neural components associated with hNR-Evs and to elucidate how hNR-EVs might act as key molecular effectors in orchestrating the spatial and temporal organization of different CNS cell types during human brain development in health and disease.

## Financial support

This work was supported by the International Society for Neurochemistry (ISN CAEN “Return Home Grant” to ALM) by the Consejo Nacional de Investigaciones Científicas y Técnicas (CONICET, PIP 11220200102807CO to CW, ALM, ACG and AC. CONICET, PIBAA 2022-2023 28820210100737CO to ALM) and the Agencia Nacional de Promoción Científica y Tecnológica (ANPCYT, PRESTAMO BID PICT 2019/00236 to ALM). MHL, MR, CW, GGG, DC, ACG, AC and ALM are investigators of CONICET.

## Acknowledgments

The authors would like to thank Dr. Sampredo, Dr. Mas and Dr. Quassollo at CEMINCO-CONICET-UNC for their support with confocal analysis. Dr. Nome and Dr. Quevedo at CIAP-INTA for their support with TEM analysis. Dr. Bazán at CIQUIBIC-CONICET-UNC for her support with DLS analysis. Dr. Eduardo Ceccarelli, Dr. Germán Rosano and Lic. Alejo Cantoia at UEM-IBR-CONICET for their support with MS analysis.

## Contributions

Conceptualization, CW, AC and ALM. Methodology, MHL., MBA, MR, LG, CW, GGG, DC, ACG and ALM Investigation, MHL, MBA, MR, LG, DC and ALM Writing - Original Draft, ALM.; Writing - Review & Editing, MR, LG, CW, GGG, ACG, AC and ALM.; Funding Acquisition, CW, ACG, AC and ALM. Resources, CW, ACG., AC and ALM.; Supervision, AC and ALM.

## Declaration of interests

The authors declare no competing interests

## FIGURE CAPTIONS

**Figure S1 |** (A) Protein levels (μg/ml media) in hiPSC-EVs and hNR-EVs. Data: mean ± SD. Unpaired t test ** < p 0.005 (2 clones: F2A112 and F2A121, 3 experiments). (B) Intersection size between top 100 EVs proteins (Vesiclepedia) and proteins identified by MS in hiPSC-EVs and hNR-EVs. (C) Abundance of transmembrane and cytosolic proteins described by MISEV 2018 as EVs markers and apolipoprotein A-1 as contaminant. (D) Western blot analysis of SDCBP (Syntenin-1) and TUBA1A levels in protein extracts from hNRs (cell lysate), hiPSC-EVs and hNR-EVs (2 clones).

**Figure S2 |** GO enrichment analysis of molecular functions (MF), cellular components (CC) and biological processes (BP) showing top 10 significantly associated terms (-log10 of adjusted p-value) in proteins detected by MS in hiPSC-EVs (A) and hNR-EVs (B). (C) hiPSCs with antibodies against PLP1 (Fire LUT), SOX2 (green) and tyrosinated tubulin (Tyr tub, magenta). Scale bar: 50 μm (20x). Scale bar 50 μm (20x). Dapi: grays.

## Notes

### Competing Interest Statement

The authors have declared no competing interest.

