## Supplementary material for "Human neural rosettes secrete bioactive extracellular vesicles enriched in neuronal and glial cellular components": Figure S1

A

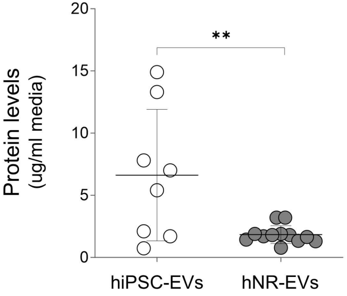


C

| Gene name | Description | hiPSC-EVs | hNR-EVs |
|---|---|---|---|
| CD9 | CD9 antigen | ND | 3.7E+06 |
| CD81 | CD81 antigen | ND | 3.5E+07 |
| CD63 | CD63 antigen | ND | 1.7E+07 |
| BSG | Basigin | ND | 1.8E+07 |
| ITGB1 | Integrin beta-1 | ND | 7.6E+07 |
| HSPA8 | Heat shock cognate 71 kDa protein | 2.2E+06 | 3.8E+08 |
| HSP90AB1 | Heat shock protein HSP 90-beta | 1.0E+07 | 2.1E+08 |
| ANXA5 | Annexin A5 | 3.3E+06 | 2.3E+07 |
| TSG101 | Tumor susceptibility gene 101 protein | ND | 4.1E+06 |
| SDCBP | Syntenin-1 | 2.8E+05 | 5.3E+07 |
| PDCD6IP | Programmed cell death 6-interacting protein | ND | 1.2E+08 |
| TUBA1B | Tubulin alpha-1B | 5.1E+06 | 2.7E+08 |
| TUBA1A | Tubulin alpha-1A chain | 4.5E+05 | 6.2E+06 |
| APOA1 | Apolipoprotein A-I | 2.3E+06 | 1.2E+06 |

B

| | EVs Top 100 (100) | hiPSC-EVs (74) | hNR--EVs (635) |
|---|---|---|---|
| Evs Top 100 (100) | | 27 | 80 |
| hiPSC-EVs (74) | 27 | | 59 |
| hNR-EVs (635) | 80 | 59 | |

D

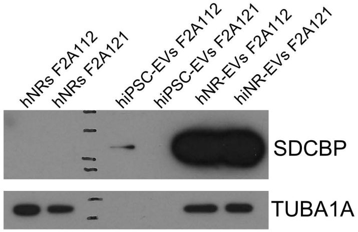
