## Supplementary figures and images for "Human neural rosettes secrete bioactive extracellular vesicles enriched in neuronal and glial cellular components"

### Figure S2

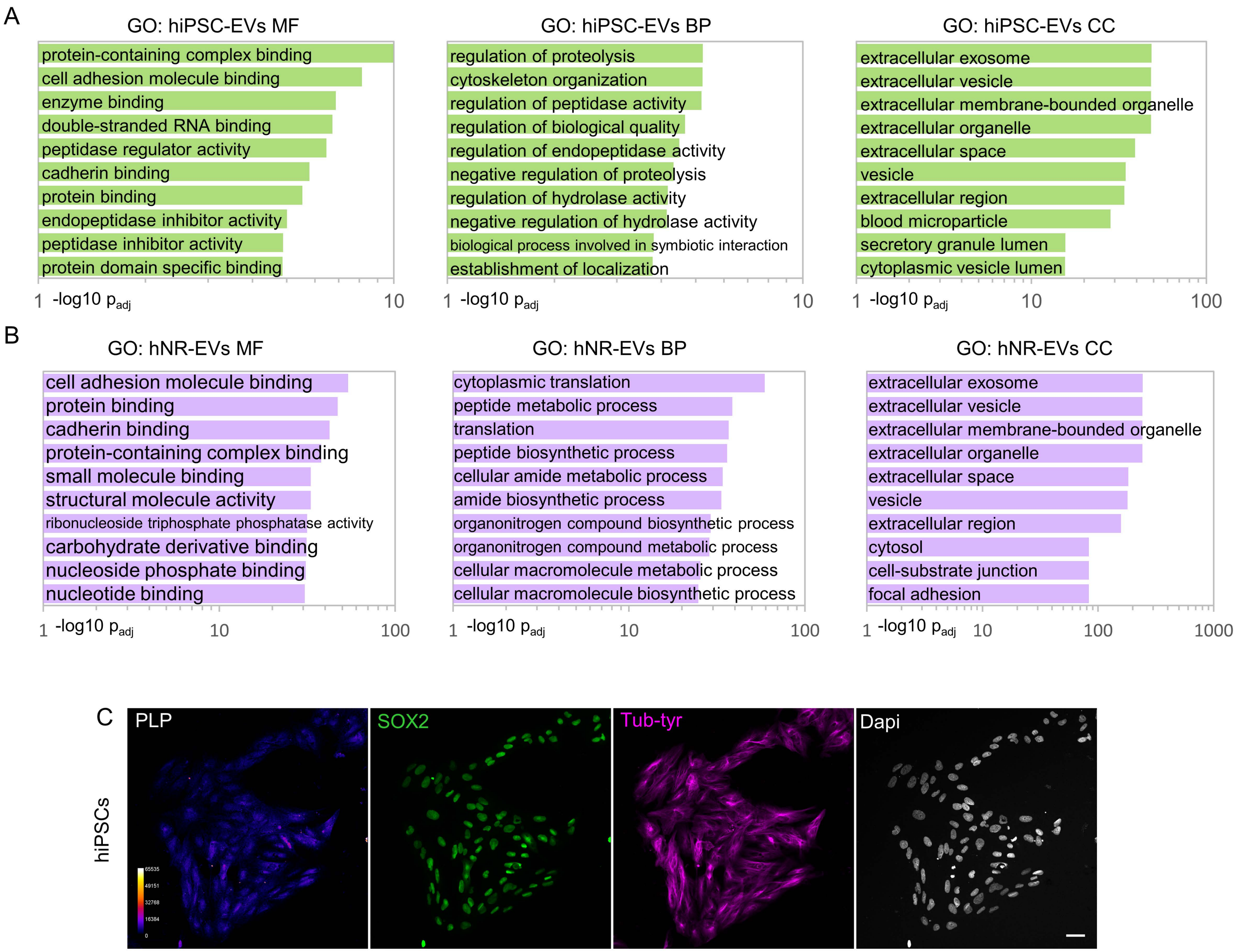
